# SVants – A long-read based method for structural variation detection in bacterial genomes

**DOI:** 10.1101/822312

**Authors:** BM Hanson, JS Johnson, SR Leopold, E Sodergren, GM Weinstock

## Abstract

**Motivation:** Mobile genetic elements (MGEs) are genetic material that can transfer between bacterial cells and move to new locations within a single bacterial genome. These elements range from several hundred to tens of thousands of bases, and are often bordered by repeat regions, which makes resolving these elements difficult with short-read sequencing data. The development and availability of long-read sequencing technologies has opened up new opportunities in the study of structural variation but there is a lack of bioinformatics tools designed to take advantage of these longer reads.

**Results:** We present an assembly-free method for identifying the location of these MGEs when compared to any reference genome (including draft genomes). Using an artificially constructed *Escherichia coli* genome containing single and tandem-repeats of a Tn9 transposon, we demonstrate the ability of SVants to accurately identify multiple insertion sites as well as count the number of repeats of this MGE. Additionally, we show that SVants accurately identifies the transposon of interest, Tn9, but does not erroneously identify existing IS1 regions present within the chromosome of the *E. coli* artificial reference.

**Availability and Implementation:** SVants is available as open-source software at https://github.com/EpiBlake/SVants

---

Mobile genetic elements (MGEs) – including plasmids, transposons, prophages, and introns – are pieces of genetic material that can rearrange within a bacterial genome as well as transfer between different bacterial cells, driving inter- and intra-species variation [1]. One of the most significant examples of the importance of MGEs in bacterial gene acquisition is the establishment, transfer, and dissemination of antimicrobial resistance genes via horizontal gene transfer (HGT) [2]. While there are a number of endogenous resistance mechanisms encoded within bacterial chromosomes, a vast repertoire of resistance mechanisms – often called the “mobile resistome” – are transferred via MGEs [3].

Historically, the study of MGEs and their insertion sites has been restricted to short-read sequencing technologies, where a number of innovative bioinformatic methods have been developed to assess structural variations in bacterial genomes [4, 5]. While these methods have helped demonstrate the importance and diversity of MGEs, limitations inherent to short reads have impeded the study of the full variety of MGEs and the structural variants they create when inserted into the genome. Specifically, short-read based methods often struggle with multiple insertion sites and repeat structures (a hallmark of many MGEs such as transposons, insertion elements, and introns) within a single genome[6, 7]. For example, IS1 is a 768-nucleotide insertion element within *Escherichia coli* K-12 MG1655 that occurs multiple times within the genome [8, 9]. As this element is longer than the longest read that short-read sequencing technologies are capable of (301 nt), it often ends up at the ends of contigs in assemblies, making it difficult to truly count the number of occurrences and the location within specific *E. coli* genomes.

Recent developments of long-read sequencing technology by Pacific Biosciences and Oxford Nanopore have enabled new ways of assessing the diversity and importance of MGEs. Oxford Nanopore in particular has shown great promise with its theoretically unlimited read length and consistent sequencing accuracy across the range of sequence lengths produced [10]. The increased read length of these platforms makes it possible to not only to identify MGEs, but also to generate long enough sequences to cover the full length of the MGE structure with additional DNA sequence on both sides. This additional DNA provides context for where MGE elements insert into the genome and bypasses all of the limitations of short-read and assembly-based methods for the identification and study of MGEs. While structural variation detection software utilizing long-read nanopore sequencing technologies have been developed for human genomics [11], assembly independent methods for the study of MGEs in bacterial genomes have not previously been described. To fill this gap and leverage the strengths of long-read sequencing, we created a tool called SVants (https://github.com/EpiBlake/SVants).

SVants is a tool to assess the presence, repeats, and insertion sites of MGEs within bacterial genomes (Figure 1a). To assess the accuracy of SVants, we created an artificial *E. coli* genome consisting of the MG1655 K-12 chromosome [12] and plasmid pO157 from enterohemorrhagic *E. coli* isolate O157:H7 Sakai [13] (Figure 1b). We then inserted a single repeat of Tn9 [14] within the chromosome, and two separate instances of Tn9 within the plasmid, one single repeat and one triple repeat. Tn9 was chosen as it is not naturally present within the MG1655 K12 genome, but MG1655 K12 contains IS1 at multiple points within its genome [12]. The presence of this IS1 element demonstrates the specificity of our tool, as we do not identify any spurious hits to chromosomal IS1 elements when looking for repeats and insertion sites of Tn9.

**Figure 1:**
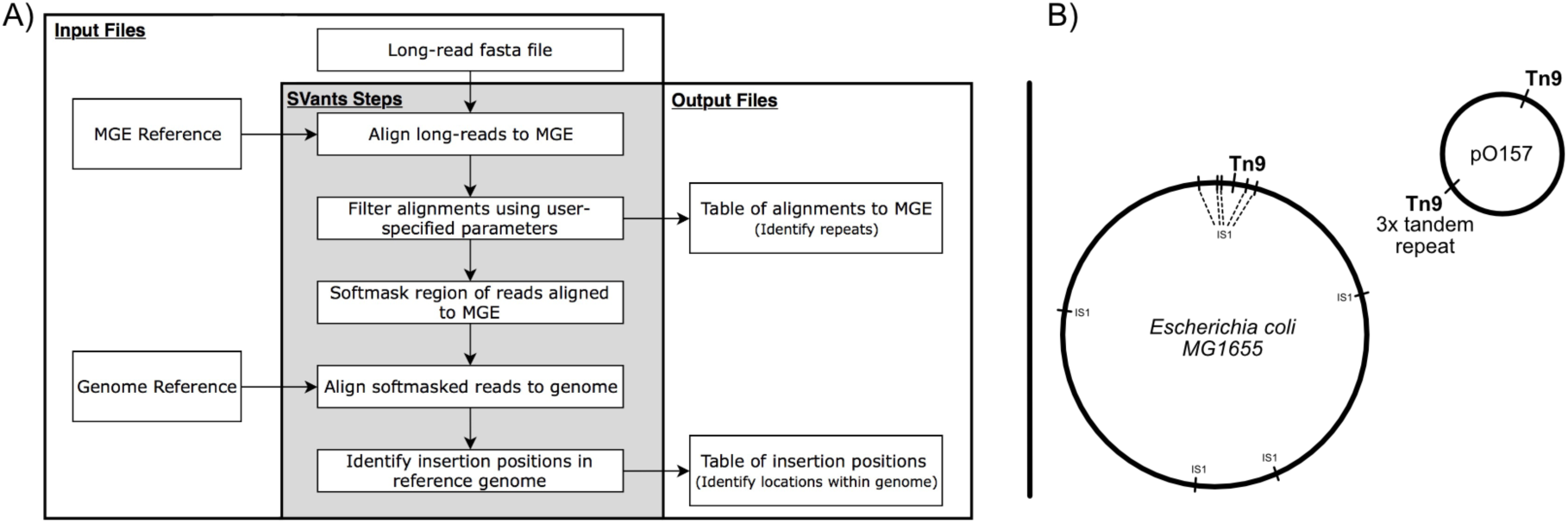
A) flow chart diagram of input files, output files, and steps completed by SVants; B) Locations of IS1 and Tn9 MGEs within the reference genome used to generate simulated Oxford Nanopore reads.

We generated 20,420 reads ranging from 35 to 100,000 bases using NanoSim [15], representing ~17x coverage of our artificial genome and roughly 1:1 coverage of the chromosome and the plasmid containing the Tn9 element. Using SVants, we were able to identify all three Tn9 insertion sites within our artificial genome; one single-repeat chromosomal insertion, one single-repeat plasmid insertion, and one triple-repeat plasmid insertion. Each of these was identified with multiple simulated reads, giving confidence to the identification of each insertion site. We were also able to show that even though there were multiple IS1 elements present within the chosen MG1655 K-12 reference genome, we only identified Tn9 insertion sites. This demonstrates the accuracy and specificity of the methods utilized within SVants. In addition to the demonstration here with an artificial *E. coli* reference, SVants has successfully been used to identify a large tandem repeat of a p1ESCUM-like plasmid within in a clinically isolated *Escherichia coli* isolate resistant to piperacillin-tazobactam [16].

Long-read sequencing technologies have opened up a new method to interrogate the diversity of MGEs and the structural variations they cause in bacterial genomes. In particular, assembly-free methods for studying these structures holds great promise. As the accuracy rate of sequencing technologies such as those from Oxford Nanopore increases, the utility of long-read sequencing for the detection and elucidation of structural variations due to MGEs will increase. Coupled with the potential for real-time analyses and antibiotic susceptibility predictions [17], detailing the complex repertoire of structural variants and MGE based mechanisms that lead to antimicrobial resistance is critical. SVants provides a new avenue for the identification of structural variation due to the presence of MGEs in bacterial genomes. This work was supported by funding from the National Institutes of Health grants 1U54 HG004968 and 1U54 DE023789 as well as funds from The Jackson Laboratory.

